# High cost of bias: Diminishing marginal returns on NIH grant funding to institutions

**DOI:** 10.1101/367847

**Authors:** Wayne P. Wahls

## Abstract

Scientific output is not a linear function of amounts of federal grant support to individual investigators. As funding per investigator increases beyond a certain point, productivity decreases. This study reports that such diminishing marginal returns also apply for National Institutes of Health (NIH) research project grant funding to institutions. Analyses of data (2006–2015) for a representative cross-section of institutions, whose amounts of funding ranged from $3 million to $440 million per year, revealed robust inverse correlations between funding (per institution, per award, per investigator) and scientific output (publication productivity and citation impact productivity). Interestingly, prestigious institutions had on average 65% higher grant application success rates and 50% larger award sizes, whereas less-prestigious institutions produced 65% more publications and had a 35% higher citation impact per dollar of funding. These findings suggest that implicit biases and social prestige mechanisms (e.g., the Matthew effect) have a powerful impact on where NIH grant dollars go and the net return on taxpayers’ investments. They support evidence-based changes in funding policy geared towards a more equitable, more diverse and more productive distribution of federal support for scientific research. Success rate/productivity metrics developed for this study provide an impartial, empirically based mechanism to do so.

## Call-Out Quotes

> “Giving the lion’s share of grant dollars to a small minority of institutions seems counterproductive and wasteful—whether or not the disparities in funding are driven by bias.”
>
> “A more egalitarian distribution of funding among institutions would yield greater collective gains for the research enterprise and the taxpayers who support it.”

## Introduction

There is strength in diversity. Diversity in scientific research includes the perspectives and creative ideas that are harnessed, the model systems and experimental tools employed, the types of investigators supported, and the regions in which research is conducted. Multiple levels of diversity increase the likelihood of scientific breakthroughs and maximize the return on taxpayers’ investments in federally sponsored research (Lorsch, 2015; Peifer, 2017a). Unfortunately, there are barriers to maximizing diversity.

A landmark study in *Science* reported that black investigators are much less likely to get their National Institutes of Health (NIH) research grant applications funded than white applicants, even after for controlling for other factors (Ginther et al., 2011). There are also large differences in success rates for investigators grouped by age (Levitt & Levitt, 2017). While there does not seem to be a gender gap for new NIH grants, female applicants have lower success rates than their male counterparts for competitive renewals (Kaatz et al., 2016; Magua et al., 2017; Pohlhaus et al., 2011). There are also large differences in success rates for investigators grouped by state (Wahls, 2016). The differences in success rates affect where federal research dollars go, contributing to heavily skewed distributions of support among all investigators. For example, just 1% of funded investigators receive about 11% of NIH research grant dollars and 10% of funded investigators get about 40% of the money (Basson et al., 2016; Collins, 2017).

One way to visualize the distribution of wealth, the magnitude of disparity and the degree of skew is through Pareto plots (**Figure 1**). The histograms (left Y-axis) display the amount of NIH research project grant funding to each bin (there are 52 bins in each plot). For example, the first bin of investigators, which contains the top-funded 1.9% of awardees, received more than twice as many dollars as the second bin (**Figure 1**, top panel). The cumulative curves (right Y-axis) display the fraction of funding that is allocated to a given bin and all higher-funded bins (i.e., those to its left). For example, the first two bins of investigators (the top-funded 3.8%) received 22% of all research dollars. Strikingly, the distributions of research dollars among institutions and states (Wahls, 2016, 2018) are even more heavily skewed than that for investigators (Basson et al., 2016; Collins, 2017). Half of all NIH research project grant dollars go to about 19% of funded investigators, 2% of funded institutions and 10% of states (**Figure 1**). The actual magnitude of disparity is even higher than depicted here because many well-qualified scientists who apply for support go unfunded. About three-quarters of applicants are denied funding each year (Rockey, 2014) and less than one in three applicants get any of their research project grant applications funded over a five-year period (Lauer, 2016c).

**Figure 1.**
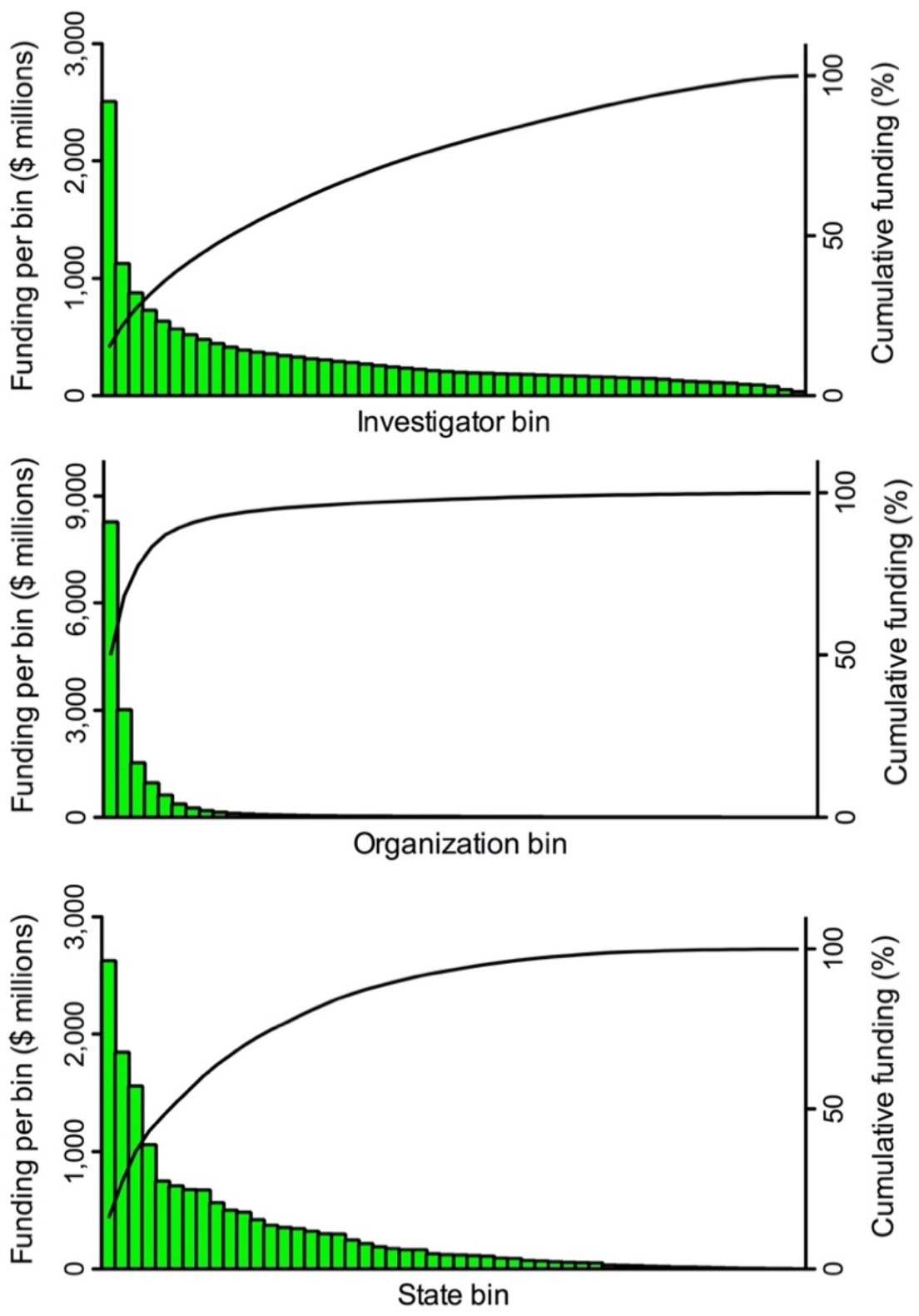
Heavily skewed distributions of NIH grant funding favor a minority and disfavor the majority. A search of the NIH RePORTER database identified 25,674 investigators who received research project grant funding in FY2015. These individuals were ranked in descending order by the amount of funding they received, and then grouped into 52 bins, each of which contained 493 investigators (the remaining, lowest-funded 38 investigators were not binned). The same process was applied for amounts of funding to 2,038 organizations (39 per bin) and to 52 states, including Washington DC and Puerto Rico (1 per bin). Pareto plots display amounts of funding (histograms, left Y axis) to each bin. Cumulative curves (right Y axis) display fraction of total funding to a given bin and all higher-funded bins (i.e., those to its left). Reproduced with permission from (Wahls, 2018) under a CC-BY 4.0 international license.

This “funding inequality has been rising since 1985, with a small segment of investigators and institutes getting an increasing proportion of funds, and investigators who start in the top funding ranks tend to stay there” (Katz & Matter, 2017). While the rich get richer, there is increasing hyper-competition elsewhere in the ranks for the remaining funds. This creates a barrier for the entry of talented young scientists into the biomedical workforce, threatening the future of the research enterprise (Carr, 2013). Similarly, the approximately 70% of awardees who hold a single NIH grant are at increased risk of losing that support, their research laboratories, and even their livelihood (Peifer, 2017a). Consequently, scientists, agency officials and organizations such as the Federation of American Societies for Experimental Biology have advocated for a more equitable distribution of funding among investigators to help sustain the biomedical research enterprise (e.g., Alberts et al., 2014; FASEB, 2015; Lorsch, 2015; Peifer, 2017a; Wahls, 2018).

Among all types of disparities in allocations of NIH funding described to date, one is preeminent—and poorly defined as to its causes and consequences. The fact that the NIH gives the majority of its extramural research project grant dollars to tiny minority (about 2%) of funded organizations (**Figure 1**) raises two fundamental, important questions. First, what factors, other than the number of applicants, contribute to the unbalanced allocations of funding among institutions? Second, are the disparities beneficial or detrimental to the national research enterprise? These questions are addressed below.

## Results

### Differences in success rates, funding rates, award sizes and funding per investigator contribute to disparity

To gain insight into potential causes of the funding disparities, funding and productivity metrics were analyzed, encompassing data over a ten-year period, for fifteen institutions whose amounts of funding ranged from about $3 million to $440 million per year (mean of values for fiscal years 2006–2015, **Supplementary Table S1**). This range extends through the first twelve bins of organizations shown in **Figure 1**, providing a broad cross section of institutions based on amounts of funding. For analyses of returns on investments, in a subsequent section of the Results, data were analyzed using continuous variable statistics. However, for part the analyses reported in this section the data were placed into groups of prestigious and less-prestigious institutions based on published rankings (Bastedo & Bowman, 2010; US News & World Report, 2016).

The first two variables examined have to do with likelihood of funding. The application-level success rate is essentially the fraction of applications that get funded in a given fiscal year, although revised applications in the same fiscal year are not counted in the denominator (Rockey, 2014). The investigator-level funding rate is the fraction of applicants that get one or more of their applications funded in a given fiscal year (Rockey, 2014). The success rates and funding rates of the institutions were obtained through a Freedom of Information Act request to the NIH (FOI case no. 46152). For fiscal years 2006 to 2015 there were about 137,000 type 1 (new) and type 2 (competing renewal) research project grant applications and the average rates for each institution in that time frame were compared. The grant application success rate for each of the prestigious institutions exceeded that for each of the less-prestigious institutions (**Figure 2A**). As a group, investigators at the prestigious institutions were, on average, 1.7-times more likely to get each grant application funded than those at the less-prestigious institutions (33.9% vs 20.5%, *p* < 0.001). Similarly, the investigator funding rate of each prestigious institution exceeded that of each less-prestigious institution, and investigators at the prestigious institutions were, on average, 1.7-times more likely to get at least one application funded each year that they applied (37.6% vs 22.4%, *p* = 0.003) (**Figure 2B**).

**Figure 2.**
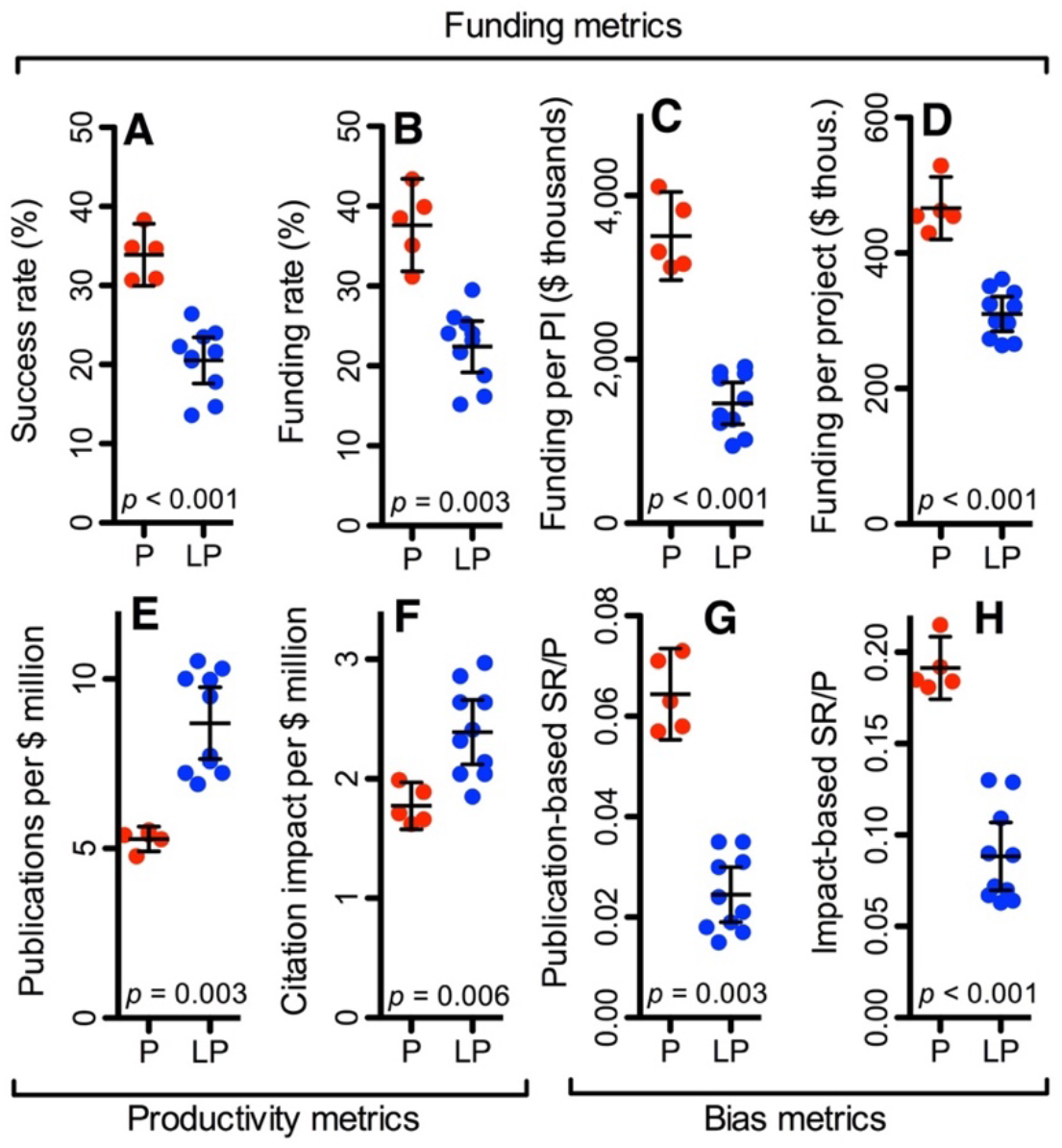
Funding allocations, productivity, and comprehensive measures of disparity. Values are for NIH research project grants, 2006–2015, grouped by prestigious (P, red) and less-prestigious (LP, blue) institutions. Funding metrics by institution are: (**A**) mean application success rate; (**B**) mean investigator funding rate; (**C**) total funding per investigator; and (**D**) mean annual funding per award. Productivity metrics are: (**E**) total publications and (**F**) total citation impact of research publications [sum of log (RCR+1)], each normalized to total funding. Differences in success rate/productivity (SR/P) ratios, using either (**G**) publication productivity or (**H**) citation impact productivity, reveal the success rate-normalized, funding amount-normalized, scientific output-normalized magnitude of disparity. Statistical values are from Mann Whitney test; lines denote mean and 95% confidence interval. Prestigious institutions: Harvard Medical School; Stanford University; Johns Hopkins University; University of California San Francisco; University of Pennsylvania. Less-prestigious institutions: Indiana University-Purdue University at Indianapolis; University of Nebraska Medical Center; University of Oklahoma Health Sciences Center; West Virginia University; University of South Dakota; Eastern Virginia Medical School, State University of New York at Buffalo; University of Mississippi Medical Center; University of North Dakota; Louisiana State University Health Sciences Center Shreveport.

The next two variables examined have to do with amounts of funding. A search of the NIH RePORTER database (US Department of Health and Human Services, 2017) identified 41,021 research project grant awards from fiscal years 2006 to 2015 (each year of funding for a project counts as an award) and these were allocated to 6,021 principal investigators. The total amount of funding to each institution over the ten years was divided by the number of investigators who received funding in one or more years to yield overall funding per investigator. The overall funding per investigator at each prestigious institution was higher than that per investigator at each less-prestigious institution (**Figure 2C**). Investigators at the prestigious institutions were awarded, on average, 2.4-times more funding than those at less-prestigious institutions ($3,508,000 vs $1,465,000, *p*< 0.001). The mean annual award size for each prestigious institution was larger than that for each less-prestigious institution, giving investigators at the prestigious institutions, on average, 1.5-times more dollars per award each year ($466,000 vs $310,000, *p* < 0.001) (**Figure 2D**).

In summary, from 2006 to 2015, each of the prestigious institutions outperformed, by every metric, each of the less-prestigious institutions in securing NIH research project grant funding.

The placement of institutions into prestigious and less-prestigious groups was part of the experimental plan, which was laid out before any data were acquired, and the assignments were based on published rankings (Bastedo & Bowman, 2010; US News & World Report, 2016). Nevertheless, these groupings could be considered arbitrary and might affect the results, so the data (**Supplemental Table S1**) were also analyzed as continuous variables without regard to prestige rank. Linear least squares regression analyses revealed robust positive correlations between success rates (R^2^ = 0.53, *p*= 0.002), funding rates (R^2^ = 0.48, *p*= 0.004), award sizes (R^2^ = 0.75, *p*< 0.001), and funding per investigator (R^2^ = 0.62, *p* < 0.001) versus the total amounts of funding to each organization.

The conclusions are straightforward. Differences in grant application success rates, investigator funding rates, annual award sizes, and funding per investigator contribute significantly to disparities in the number of research project grant dollars allocated to institutions. Moreover, the impacts of the differences in success rates (**Figure 2A**) and award sizes (**Figure 2D**) are multiplicative, giving the prestigious institutions about 240% more dollars of funding per investigator (**Figure 2C**). In short, differences in likelihood of funding and award sizes are proximate causes of the heavily skewed distribution of funding among institutions (**Figure 1**). Consequences of these imbalances are documented in subsequent sections of the Results and are described in the Discussion.

### Less-prestigious institutions produce greater returns on investments

The disparities in allocations of funding by the NIH might be justified if the prestigious institutions were of greater value to the national research enterprise than the less-prestigious institutions. To see if this is the case, I examined two variables for their primary scientific outputs, which are funding-normalized publication productivity and the citation impacts of those publications.

There were 41,021 research project grant awards from 2006 to 2015. The project numbers for awards to each institution were used to search the PubMed database (US National Library of Medicine and National Institutes of Health, 2017), which identified 95,035 scientific publications (based on their unique PMIDs) that were supported by those projects from 2006 to 2015. The total number of project-associated publications of each institution was divided by total funding to yield publication productivity. Each of the less-prestigious institutions produced more scientific publications per dollar of research project grant funding than each of the prestigious institutions (**Figure 2E**). They were, on average, 65% more productive (8.7 vs 5.3 publications per million dollars of funding, *p* = 0.003). Of course, it is possible that the scientific impact of publications might differ between institutions.

To gain insight into this possibility, the relative citation ratio (RCR) (Hutchins et al., 2016) was compiled for each grant-supported research article during the survey period. Citations to reviews, editorials, and other non-research article types were excluded from analysis. The RCR value, which is being used by the NIH to assess portfolio performance and to guide funding decisions (e.g., Lauer, 2016a, 2016b, 2016d, 2017), is a time-normalized, field-normalized metric for citation impact (Hutchins et al., 2016). These normalizations allow one to compare, in an appropriately weighted fashion, the impact factors for articles published at different times in the survey period. Since article-level citation impact factors follow a log-normal distribution (Eom & Fortunato, 2011; Hutchins et al., 2016; Stringer et al., 2008), RCR (+ 1) values were log-transformed (e.g., Kaltman et al., 2014). The sum of log-RCR values for each institution was normalized to total funding, which provides a measure of productivity based on the citation impact of publications. All but one of the less-prestigious institutions outperformed each of the prestigious institutions, and as a group they had a 35% higher productivity (**Figure 2F**, *p* = 0.006).

In summary, from 2006 to 2015, the overall, funding-normalized productivity of the less-prestigious institutions was greater than (35% based on citation impact) or substantially greater than (65% based on publication rate) that of the prestigious institutions. I conclude that the scientific output-based of value of these institutions to the national research enterprise does not justify the strong disparities in allocations of funding (significant differences in success rates, funding rates, award sizes, and funding per investigator) between the prestigious and less-prestigious institutions.

It should be emphasized that the differences in productivity do not necessarily mean that investigators at the less-prestigious institutions are “better scientists” or are “more meritorious” than those at the prestigious institutions. Reasons for this are documented in a subsequent section of the Results and are described in the Discussion.

### A more comprehensive measure for the magnitude of disparity

Previous studies of funding disparities have focused primarily on differences in grant application success rates (e.g., Ginther et al., 2011; Kaatz et al., 2016). However, results of this study and those recently reported elsewhere (Murray et al., 2016; Wahls, 2016) show that there are also disparities in amounts of funding per award. When investigators who are in a group that is disadvantaged by lower success rates do get their applications funded, they often receive substantially less money per award (e.g., **Figure 2D**). Moreover, there can be substantial differences in productivity between groups (e.g., **Figure 2E-2F**), which is germane to whether differences in success rates and award sizes are warranted. These various factors can be evaluated simultaneously by using the SR/P value, which is success rate divided by productivity. Differences in SR/P values for investigators grouped in any way that is desired (e.g., by race, gender, age, institution or state) and using any measure of productivity that is desired (e.g., publication rate or citation impact per unit of funding), reveal the success rate-normalized, funding amount-normalized, scientific output-normalized magnitude of funding disparities.

For all four of the different ways that the data were analyzed, the SR/P value (and a related metric, below) of each prestigious institution exceeded that of each less-prestigious institution (**Figure 2G-2H** and **Supplementary Table S1**). When publications were used as the basis for productivity, the mean SR/P value of the prestigious institutions was 2.6-fold higher than that for the less-prestigious institutions (**Figure 2G**, *p* = 0.003). When citation impact values were used to gauge productivity, there was a 2.2-fold difference between groups (**Figure 2H**, *p*< 0.001). Substituting per investigator funding rates (FR) for per application success rates (SR) produced essentially identical results, with intergroup FR/P quotients of 2.7 (*p* = 0.003) and 2.2 (*p* = 0.003), respectively (**Supplementary Table S1**). The fact that four distinct approaches yielded concordant results (mean of 2.41 ± 0.27 standard deviation) suggests that SR/P and FR/P metrics developed for this study provide robust measures for the magnitude of disparity.

### Inverse correlations between amounts of funding and productivity

To gain insight into consequences of the funding disparities, publication-based and citation impact-based productivity values were analyzed as a function of total funding, mean annual funding per award, and funding per principal investigator at each institution (**Figure 3**). For each of these six analyses, linear regression statistics revealed a robust inverse correlation between amounts of funding and productivity (R^2^ = 0.53 to R^2^ = 0.78; *p* < 0.001 to *p*= 0.003). I conclude that there are diminishing marginal returns on allocations of NIH research project grant dollars among these institutions, as reported for amounts of NIH funding among individual grants (e.g., Lauer, 2016a, 2016b), investigators (e.g, Basson et al., 2016; Lorsch, 2015), and quartiles of states (Wahls, 2016). The causes of such diminishing marginal returns, their impacts on the national research enterprise, and implications for funding policy are presented in the Discussion section.

**Figure 3.**
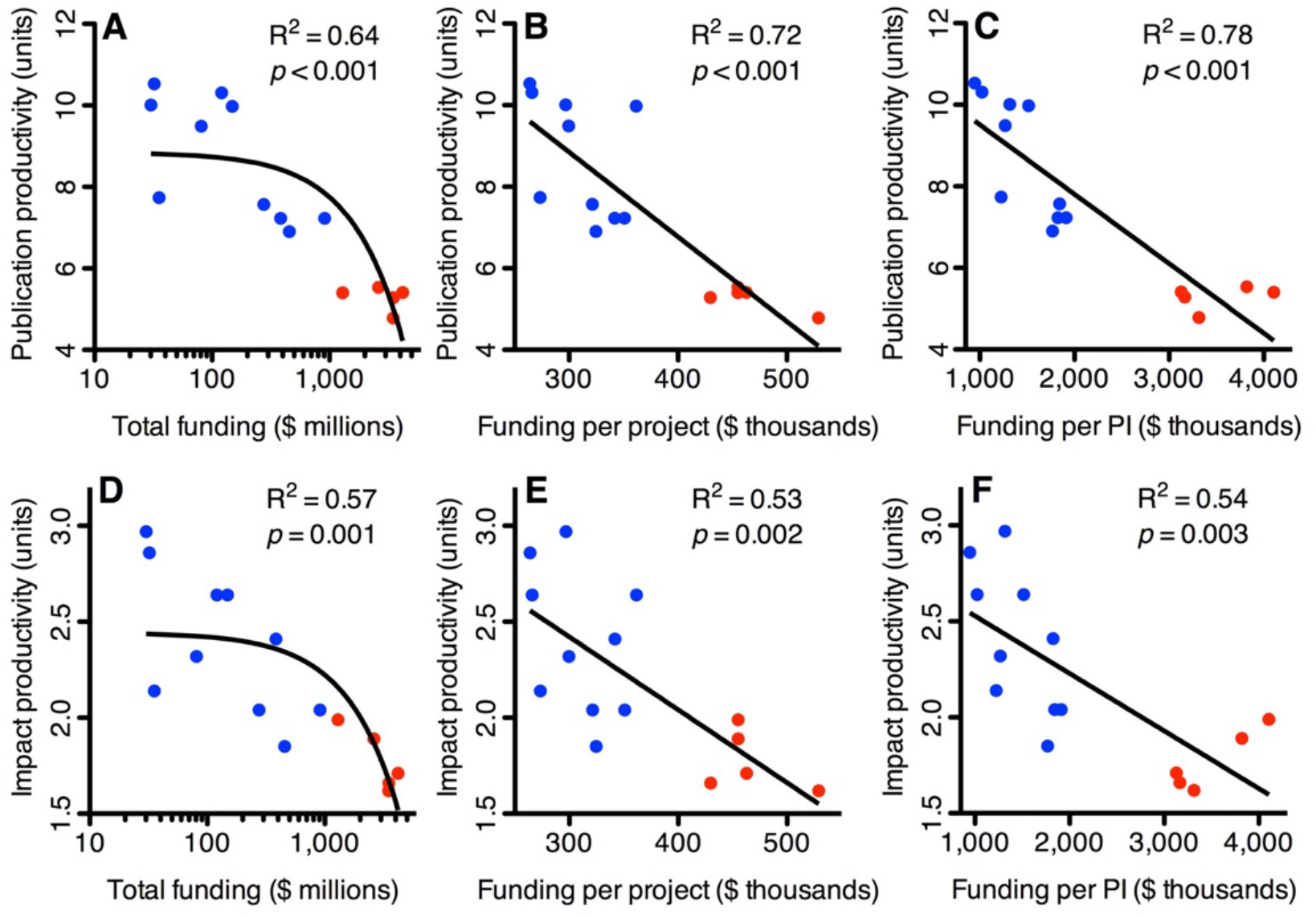
Effects of funding disparities on productivity. (**A-C**) Publication productivity and (**D-F**) citation impact productivity are plotted as a function of (**A, D**) total funding; (**B, E**) annual funding per project; and (**C, F**) funding per investigator at each institution. Data for prestigious and less-prestigious institutions are shown in red and blue, respectively. Lines and statistical values are from linear regression; curvatures in panels **A** and **D** are due to plotting total funding on a log scale.

### Generalizability of the findings

The analyses encompassed institutions whose amounts of funding ranged from $3 million to $440 million per year and the conclusions are based on statistically significant differences in data from more than 100,000 research project grant applications, 40,000 awards, and 95,000 publications acknowledging support from those grants over a ten-year period. Inspection of the literature revealed that the differences in grant application success rates reported here for a subset of institutions (65% difference between groups) are virtually identical to those reported for all institutions placed in groups by their amounts of grant funding (Eblen et al., 2016) and, in another study, for all institutions grouped by size (Murray et al., 2016). Similarly, the differences in award sizes reported here are like those reported for all institutions (Murray et al., 2016). The findings of this study, using a cross section of institutions whose amounts of funding cover a broad (about 150-fold) range, can thus be considered representative of the broader population of institutions.

## Discussion

There are three key findings described in this study. First, allocations of NIH research project grant funding to institutions are extremely skewed, favoring a tiny minority and disfavoring the vast majority (**Figure 1**). Second, differences in grant application success rates and award sizes contribute to these disparities (**Figure 2A,2D**). The impacts of differences in success rates and award sizes are multiplicative, giving the favored institutions about 240% more dollars per investigator (**Figure 2C**). Third, the scientific productivity of the disfavored institutions exceeds that of the favored institutions (**Figure 2E-2F**) and there are robust inverse correlations between funding (total, per award, per investigator) and productivity (**Figure 3**). These findings provide important new insight into causes and consequences of disparities in federal funding for scientific research, and they support evidence-based changes in funding policy.

### Funding allocations are biased by institution

The extreme disparities in NIH funding to institutions (e.g., 1% of funded organizations get about 34% of the dollars), which favor a tiny minority and disfavor the vast majority (**Figure 1**), are not matched by extreme differences in distributions of talent. For example, a congressionally mandated study found that the talent to carry out research resides throughout the United States (National Academies, 2013). All institutions have access to a surplus of highly trained investigators and supporting scientists (Alberts et al., 2014; Carr, 2013) and the value of an investigator to the nation’s research enterprise is largely independent of institutional affiliation (Deville et al., 2014). Moreover, this study revealed that large differences in grant application success rates and award sizes among institutions are discordant with their productivity-based value to the national research enterprise (**Figure 2**). It thus appears that the NIH funding process is biased by institution, as has been reported for funding by the Natural Sciences and Engineering Research Council of Canada (Murray et al., 2016).

### Subconscious bias and social prestige mechanisms

It seems unlikely that grant reviewers and NIH officials at-large are overtly biased, so what are potential sources of bias and how could they possibly have such a strong impact on allocations of funding among institutions?

Most bias is subconscious and these pervasive, implicit biases even affect the actions of individuals who are not overtly biased (Lai et al., 2013; Staats et al., 2016). Our actions are also strongly affected by social prestige mechanisms that encompass non-meritocratic factors such as the wealth, reputation and selectivity of institutions (Bastedo & Bowman, 2010; Burris, 2004; Clauset et al., 2001). The preferential allocation of NIH funding to prestigious institutions (**Figure 2**), despite their lower productivity, is an excellent example of the Matthew effect (a type of bias/social prestige mechanism) (Merton, 1968; Perc, 2014) in action. As another example, manuscripts are more frequently accepted for publication when they come from prestigious institutions than from less-prestigious institutions, and the acceptance rate gap closes when author identity and institutional affiliation are withheld from the reviewers (Tomkins et al., 2017). We are hard-wired, biologically, to make conscious and subconscious distinctions between groups of people and those distinctions, however unjustified they might be, can affect allocations of funding.

A little bias goes a long way. Even small differences in reviewers’ scores for preferred and non-preferred applicants produce large differences in grant application success rates (Day, 2015). There are at least four distinct steps of the funding process, involving both scientific merit review (peer review) and administrative funding decisions, at which bias can occur (**Figure 4**). Consequently, the effects of even minor, subconscious biases at each step can multiply exponentially through successive steps of the process. Their net impact at population scale can be inferred by measuring differences in SR/P values, which take into account differences in likelihood of funding, amounts of funding, and scientific output between investigators grouped in any way desired. Four different permutations of this metric yielded similar results (**Figure 2G-2H, Supplementary Table S1**) for the magnitude of disparity between the groups of prestigious and less-prestigious institutions analyzed (mean of 2.41 ± 0.27 standard deviation). The SR/P metric thus provides a potentially useful benchmark for ameliorating disparities and, as described below, for optimizing the efficiency with which research dollars are expended.

**Figure 4.**
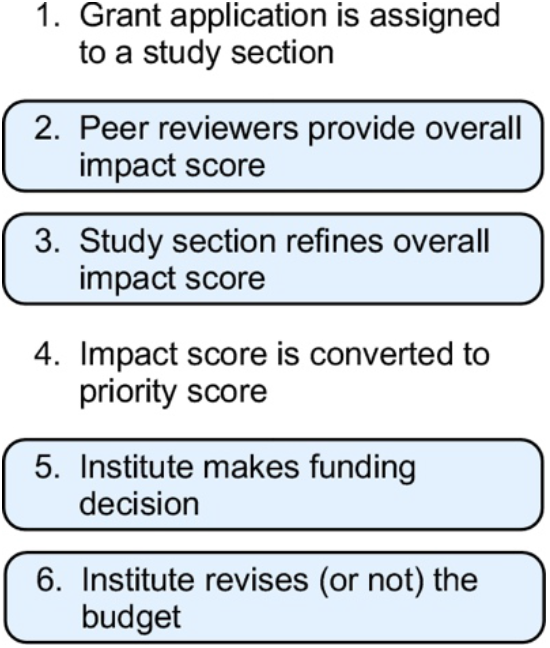
Multiple opportunities for bias. The impacts of even minor, subconscious biases at individual steps of the funding process (shaded) can multiply exponentially [effects of (bias 1) ´ (bias 2) ´ (bias 3) ´ (bias 4) = (net impact of bias)].

### Disparities in funding affect the return on taxpayers’ investments

The principle that unbalanced allocations of grant funding yield diminishing marginal returns (incremental output for each additional dollar of funding) has been documented extensively at the level of investigators (e.g., Basson et al., 2016; Berg, 2010; Cook et al., 2015; Doyle et al., 2015; Fortin & Currie, 2013; Lauer, 2016a, 2016b; Lorsch, 2015; Mongeon et al., 2016). It stems from the fact that individual investigators each have a finite capacity to carry out grant-related duties and their productivity declines when their amounts of funding exceed those capacity limits (Alberts, 1985). At population scale these diminishing marginal returns, which are a direct consequence of giving a disproportionately large share of grant funding to a minority of investigators, have profound impacts on how efficiently research dollars are being expended. For example, analyses of National Institute of General Medical Sciences (NIGMS) award data revealed that funding for one R01 grant to an investigator produces, on average, about five scientific publications in the funding period, whereas the same amount of funding for a third R01 grant yields only about one additional publication (Lorsch, 2015). As another example, based on NIH-wide funding data and citation impact factors (median RCR values), marginal returns for investigators with $400,000 of annual research project grant funding are about five-times greater than those for investigators with a million dollars of funding (Lauer et al., 2017). The diminishing marginal returns persist even when investigator award data are parsed by NIH institute, for “elite” investigators, and by human versus non-human model systems (Lauer et al., 2017).

This study revealed that diminishing marginal returns also apply at the level of institutions (**Figure 3**). The ramifications of this finding are like those for returns on investments at the level of investigators. Because the NIH gives half of all research project grant dollars to about 2% of supported institutions (the very well-funded ones) (**Figure 1**) and very well-funded institutions tend to be considerably less productive than more modestly funded institutions (**Figure 2E-2F, Figure 3**), the unbalance allocations have profound implications for the efficiency with which research dollars are being expended. Giving the lion’s share of grant dollars to a small minority of institutions seems counterproductive and wasteful—whether or not the disparities in funding are driven by bias. As is the case for the distribution of research dollars among individual investigators (Lorsch, 2015; Mongeon et al., 2016; Peifer, 2017a, 2017b; Wahls, 2017, 2018), a more egalitarian distribution of funding among institutions would yield greater collective gains for the research enterprise and the taxpayers who support it.

### SR/P values provide impartial way to reduce disparity and increase return on investments

To effectively reduce systemic disparities in allocations of funding (e.g., **Figure 1**), the NIH would have to close gaps in grant application success rates and award sizes for investigators grouped by race (Ginther et al., 2011), gender (Kaatz et al., 2016; Magua et al., 2017; Pohlhaus et al., 2011), age (Levitt & Levitt, 2017), institution (this study) and state (Wahls, 2016). The mechanism for remediation would also have to address the impacts of diminishing marginal returns (e.g., **Figure 3**) and, furthermore, must do so in proportion to their variable magnitude. Overall, the process would have to strike a balance between three fundamental needs: First, ensure that investigators at-large are allowed to compete on equal footing for grants and grant dollars. Second, accommodate the possibility that some groups of investigators might be of greater value to the research enterprise than other groups. Third, maximize the net return on taxpayers’ investments. The SR/P metrics developed for this study provide a straightforward and impartial way to satisfy, simultaneously, these three fundamental needs.

The differences in SR/P values between institutions (**Figure 2G-2H**) encompass the impacts of diminishing marginal returns on scientific output (productivity) as well as controllable factors (differences in success rates and award sizes) that contribute to the diminishing marginal returns. Thus, SR/P values provide useful parameters with which to optimize the net return on taxpayers’ investments. To do so, the NIH would adjust success rates and award sizes to the extent that is necessary to establish parity or near parity of SR/P values between institutions. Success rates and award sizes could still vary between institutions (according to their productivity-based merit), up to but not exceeding the point at which their SR/P values depart from the target range. This approach would treat systematically and proportionately the proximate causes of institutional funding disparities and their deleterious impacts on net productivity of the research enterprise. Moreover, because SR/P values can be derived for investigators grouped in any way desired, the proposed mechanism is of broad utility for addressing imbalances in funding allocations and net productivity among populations of investigators grouped in other ways (e.g., by race, gender, age and state).

### Summary and implications for funding policy

In conclusion, this study and others (e.g., Basson et al., 2016; Berg, 2010; Cook et al., 2015; Doyle et al., 2015; Fortin & Currie, 2013; Lauer, 2016a, 2016b; Lauer et al., 2017; Lorsch, 2015; Mongeon et al., 2016; Wahls, 2016) support evidence-based changes in funding policy geared towards a more equitable, more diverse and more productive distribution of federal support for scientific research. A wealth of data, such as differences in SR/P values (**Figure 2**) and returns on taxpayers’ investments (**Figure 3**), document unambiguously the need for such changes— and provide empirical benchmarks for remediation.

## Methods

### Data sets

Data on funding and productivity by institution for FY2006 to FY2015 are provided in **Supplementary Table S1**. The institutions were selected from published rankings (US News & World Report, 2016). Five institutions were from the top of the list and the remainder were selected at random from mid-ranked, low-ranked, rank not posted, and unranked regions of the list to provide a cross-section of institutions. Data on research project grant application success rates and investigator funding rates of each institution for FY2006 to FY2015 were obtained from the NIH Office of Extramural Research (Tables #96–17–1 and #96–17–2; in response to FOI case no. 46152). The means of all type 1 (new) and type 2 (competing renewal) applications from FY2006 to FY2015 were determined. Data on total number of research project grant awards, investigators, and funding from FY2006 to FY2015 were obtained by searching the NIH RePORTER database (US Department of Health and Human Services, 2017). Search parameters were institution (using organization-specific DUNS numbers), fiscal year (2006–2015), and funding mechanism (research project grants). A list of each institution’s grant numbers from the RePORTER search was constructed and was used to search the PubMed database (US National Library of Medicine and National Institutes of Health, 2017) for the number of grant-supported publications from 2006 to 2015. The list of PMIDs for grant-supported publications of each institution was used to search the iCite database (Hutchins et al., 2016) to obtain the relative citation ratio of each publication. Additional data sets were derived algebraically as described in the Results and **Supplementary Table S1**.

### Statistical tests

Grouped data sets were analyzed using the Mann Whitney test; continuous variable data sets were analyzed using linear least squares regression; analyses were conducted in Prism (GraphPad Software, Inc., La Jolla, CA, USA).

### Data availability

All relevant data are contained in the manuscript and its Supplementary Information file. Additional datasets (e.g., raw results from searches of NIH RePORTER and PubMed) are available from the corresponding author upon request.

## Acknowledgments

I thank the NIH Office of Extramural Research for providing data and G. Baldini, T. Chambers, L. Cornett, M. Davidson, and F. Kilic for comments on the manuscript.

## Supplementary Information

**Table S1.** Prestige rank, funding, publication, citation impact, and funding-normalized productivity data by institution (2006–2015).

| Rank <sup>1</sup> | DUNS No. <sup>2</sup> | Organization | Success Rate <sup>3</sup> | Funding Rate <sup>3</sup> | Projects <sup>4</sup> | Funding <sup>4</sup> | Funded Pls <sup>4</sup> | Funding / Projects |
| --- | --- | --- | --- | --- | --- | --- | --- | --- |
| 1 | 047006379 | Harvard Medical School | 38.3% | 43.4% | 2,634 | \$1,297,809,360 | 316 | \$455,023 |
| 2 | 009214214 | Stanford University | 34.7% | 39.9% | 5,771 | \$2,625,937,078 | 687 | \$455,022 |
| 3 (tie) | 001910777 | Johns Hopkins University | 30.9% | 31.2% | 9,534 | \$4,412,893,242 | 1,411 | \$462,859 |
| 3 (tie) | 094878337 | University of California, San Francisco | 34.8% | 38.5% | 7,453 | \$3,493,270,490 | 1,054 | \$529,085 |
| 3 (tie) | 042250712 | University of Pennsylvania | 30.7% | 35.1% | 8,159 | \$3,506,420,297 | 1,103 | \$429,761 |
| 47 | 603007902 | Indiana University-Purdue University at Indianapolis | 22.3% | 25.3% | 2,597 | \$911,515,992 | 477 | \$350,998 |
| 63 | 168559177 | University of Nebraska Medical Center | 21.6% | 24.1% | 1,122 | \$383,461,292 | 210 | \$341,766 |
| 72 | 878648294 | University of Oklahoma Health Sciences Center | 26.4% | 29.5% | 860 | \$276,457,268 | 150 | \$321,462 |
| 86 | 191510239 | West Virginia University | 17.8% | 18.8% | 454 | \$120,706,995 | 118 | \$265,874 |
| 88 | 929930808 | University of South Dakota | 20.9% | 21.7% | 102 | \$30,274,425 | 23 | \$296,808 |
| RNP | 058625146 | Eastern Virginia Medical School | 13.6% | 15.2% | 130 | \$35,552,814 | 29 | \$273,483 |
| RNP | 038633251 | State University of New York at Buffalo | 24.0% | 26.1% | 1,401 | \$454,795,551 | 257 | \$324,622 |
| UR | 928824473 | University of Mississippi Medical Center | 23.5% | 24.1% | 411 | \$148,645,596 | 98 | \$361,668 |
| UR | 102280781 | University of North Dakota | 20.5% | 23.3% | 122 | \$32,202,013 | 34 | \$263,951 |
| UR | 095439774 | Louisiana State University Health Sciences Center Shreveport | 14.7% | 16.2% | 271 | \$81,206,676 | 64 | \$299,656 |

<sup>1</sup> Rank order from 2016 US News & World Report list of "Best Medical Schools: Research" (RNP, rank not posted; UR, unranked).
<sup>2</sup> Organization-specific DUNS numbers were used for searches of NIH RePORTER.
<sup>3</sup> Rates are means of all type 1 (new) and type 2 (renewal) RPG applications, FY2006-FY2015, from NIH OER Tables #96-17-1 and #96-17-2.
<sup>4</sup> Totals for FY2006-FY2015 are from searches of NIH RePORTER conducted from 06/17/2016 to 06/21/2016.
<sup>5</sup> Grant-supported publications in 2006-2015 from searches of PubMed on 06/24/2016.

| Funding / Pls | Publications <sup>5</sup> | Pubs / \$M<br>Funding | Citation<br>Impact <sup>6</sup> | Impact / \$M<br>Funding | Publication-based SR / P <sup>7</sup> | Impact-based<br>SR / P <sup>7</sup> | Publication-based FR / P <sup>7</sup> | Impact-based<br>FR / P <sup>7</sup> |
| --- | --- | --- | --- | --- | --- | --- | --- | --- |
| \$4,106,992 | 7,004 | 5.40 | 2577 | 1.99 | 0.071 | 0.192 | 0.080 | 0.218 |
| \$3,822,325 | 14,541 | 5.54 | 4971 | 1.89 | 0.063 | 0.184 | 0.072 | 0.211 |
| \$3,127,493 | 23,884 | 5.41 | 7552 | 1.71 | 0.057 | 0.181 | 0.058 | 0.182 |
| \$3,314,298 | 16,692 | 4.78 | 5668 | 1.62 | 0.073 | 0.215 | 0.081 | 0.238 |
| \$3,167,199 | 18,333 | 5.28 | 5832 | 1.66 | 0.058 | 0.185 | 0.066 | 0.211 |
| \$1,910,935 | 6,588 | 7.23 | 1861 | 2.04 | 0.031 | 0.109 | 0.035 | 0.124 |
| \$1,826,006 | 2,844 | 7.23 | 924 | 2.41 | 0.030 | 0.090 | 0.033 | 0.100 |
| \$1,843,048 | 2,092 | 7.57 | 565 | 2.04 | 0.035 | 0.129 | 0.039 | 0.145 |
| \$1,022,941 | 1,244 | 10.31 | 319 | 2.64 | 0.017 | 0.067 | 0.018 | 0.071 |
| \$1,316,279 | 303 | 10.01 | 90 | 2.97 | 0.021 | 0.070 | 0.022 | 0.073 |
| \$1,225,959 | 275 | 7.74 | 76 | 2.14 | 0.018 | 0.064 | 0.020 | 0.071 |
| \$1,769,632 | 3,136 | 6.90 | 840 | 1.85 | 0.035 | 0.130 | 0.038 | 0.141 |
| \$1,516,792 | 1,484 | 9.98 | 392 | 2.64 | 0.024 | 0.089 | 0.024 | 0.091 |
| \$947,118 | 339 | 10.53 | 92 | 2.86 | 0.019 | 0.072 | 0.022 | 0.081 |
| \$1,268,854 | 771 | 9.49 | 188 | 2.32 | 0.015 | 0.063 | 0.017 | 0.070 |

